# First report of the Eocene bivalve *Schedocardia* (Mollusca, Cardiidae) from Cuba

**DOI:** 10.1101/2020.02.03.932756

**Authors:** Johanset Orihuela, Yasmani Ceballos Izquierdo, Roger W. Portell

## Abstract

Herein we provide the first report of the cardiid bivalve *Schedocardia* from Cuba. The single, partial, valve external mold was derived from the Madruga Formation which is characterized by a richly diverse marine fauna including echinoderms, brachiopods, benthic and planktonic foraminiferans, but from which bivalves were not previously reported. The unit is considered late Paleocene in age (Thanetian), but the presence of *Schedocardia* supports a possible age extension of the formation into the early Eocene (Ypresian). Moreover, we provide a reconsideration of the historical factors that affected the accepted type locality of the outcrop, which allows for an alternative interpretation of the fauna found therein.

**RESUMEN:** Se reporta por primera vez el género bivalvo *Schedocardia* de la Formación Madruga, para el registro fósil de Cuba. De esta formación se ha reportado una rica fauna de invertebrados incluyendo equinodermos, braquiópodos y, especialmente de foraminíferos, pero no moluscos bivalvos. Se ha considerado que la Formación Madruga se depositó durante el Paleoceno tardío (Thanetiano), no obstante, la presencia de *Schedocardia* apoya la extensión de edad de la formación hasta el Eoceno temprano (Ypresiano). Además, se consideran factores antropogénicos que han afectado la localidad tipo de la formación.

## INTRODUCTION

The Paleocene of Cuba is represented in localized outcrops throughout the island, but a characteristic one is located near the town of Madruga, in the central-eastern part of the Mayabeque Province, where a complex of ophiolites, Cretaceous volcanic rocks, and latest Cretaceous, Paleogene and Neogene age sedimentary formations crop out (Furrazola-Bermúdez et al., 1964; Albear et al., 1985).

This region has been the subject of paleontological investigations since the early-middle 20^th^ century, as part of oil and mineral prospecting by North American companies (De Golyer, 1918; Lewis, 1932a, b; Palmer, 1932; Wright and Sweet, 1924). Based on field trips made between 1929 and 1946 around Cuba, Palmer (1948) compiled a list of fossil-bearing localities, including more than 100 sites for the town of Madruga. In his publication, Palmer mentioned a locality previously identified by Lewis (1932a) as “*Cut under railroad bridge two kilometers west of Madruga at Central San Antonio*” and cataloged it as a notable locality (Loc. 757 in Palmer, 1948). Since Lewis (1932a) had not selected a holostratotype (type section) nor did he record its faunal content, Bermúdez (1950) erected the roadcut under the railroad bridge as the type locality for the Madruga Formation (Loc. Bermúdez Sta. 76b) and assigned it a Paleocene age.

This locality was originally described as “Madruga marls” by Lewis (1932a), who erroneously considered it of Late Cretaceous age. Its fossil fauna was reported later by Palmer (1934), Palmer (1932, 1948), Cushman and Bermúdez (1948a, b, 1949), Bermúdez (1938, 1950), Cooper (1955, 1979), Sachs (1959), and Kier (1984). They had found mostly Cretaceous and Paleocene large foraminifers, but also assemblages consisting of small benthic and planktonic species including several index taxa (Palmer, 1934; Cushman and Bermúdez, 1948a, b, 1949; Bermúdez, 1950; Furrazola-Bermúdez et al., 1964; Franco-Álvarez et al., 1992). Interestingly, brachiopods, scaphopods, and echinoid fragments were also reported. The brachiopods collected by Palmer, deposited in the USNM (United States National Museum, Washington D.C.), were later described and figured by Cooper (1955, 1979). Although these works provide a detailed account of the foraminifer faunule from the Madruga outcrop, bivalves had not been reported, and the echinoids and scaphopods still need investigation.

Recent fieldwork at this locality yielded several interesting, well-preserved fossil bivalve molds, representing a new addition to the fauna of the Madruga Formation undetected in over 80 years of collecting history at the site. Furthermore, our new observations contribute to the reinterpretation of the age and environment of deposition of the Madruga Formation.

## MATERIALS AND METHODS

### Locality

The Madruga Formation type section and collecting site of the bivalves herein reported is located at a highway cut near the Boris Luis Santa Coloma Sugar Plantation, outskirts of the town of Madruga, Mayabeque Province: GPS coordinates Lat. 22.906787°, Long. -81.872793°, datum WGS 84, altitude 159 m above modern sea level (taken from Google Earth). This is the same location, below the railroad bridge at the Carretera Central west of Madruga town, reported by Palmer (1948) as Loc. 757, and Loc. Sta. 76b in Bermúdez (1950). This section is the current lectostratotype of the Madruga Formation (Albear et al., 1985; Franco-Álvarez et al., 1992). The outcrop section discussed is ∼4 m above the highway level, on each side. The bivalves herein described were collected at a +1 m and +2.5 m (Fig. 1). This is nearby the original localities visited by Palmer and Bermúdez, but the exact locality of their collecting stations is unknown because they did not indicate them.

**Figure 1:**
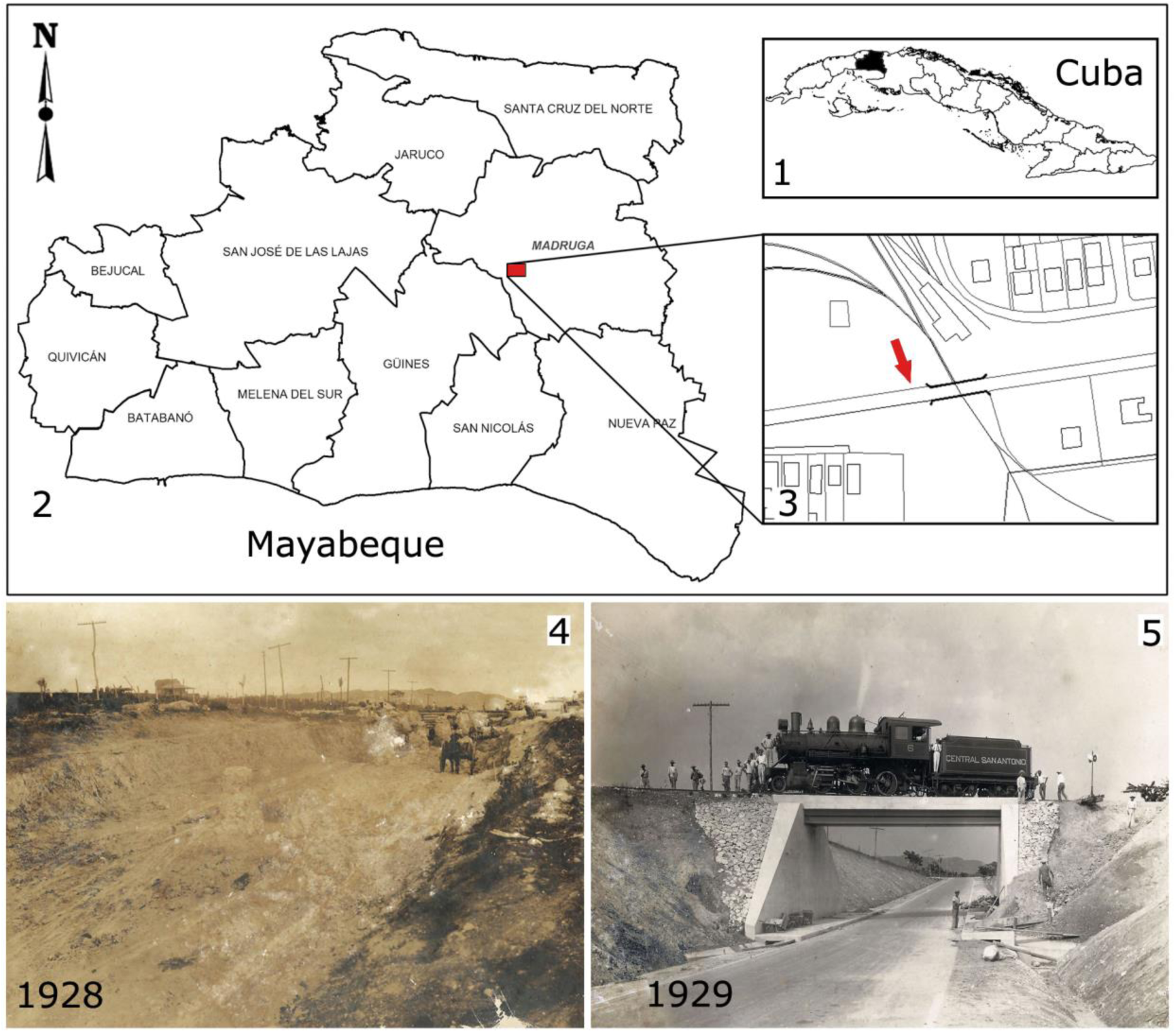
Collecting locality of the Madruga Formation, near the city of Madruga, Province of Mayabeque, Cuba. 1.1 Mayabeque province, 1.2 Madruga municipality, and 1.3 Madruga Formation type locality indicated by the red arrow near the railroad bridge. Historical photographs of the construction of the road under the railroad bridge, where the exposed outcrops of the Madruga Formation can be seen during construction (1.4) and after (1.5).

The specimens were collected from loose fragments found within the conglomerates that make up the stratigraphic column of the formation at the referenced locality (Fig. 1). Their lithology, coloration, and position within the column indicate they are loose clastics of the Madruga Formation, and not the Late Cretaceous bed that lies discordantly below.

### Geological setting: Lithology and Deposition environment

The Madruga section has been described as brown or reddish-brown calcareous well-stratified, cemented, greywacke sandstones, sandy-shales, and shales with polymictic conglomerate intercalations. This formation lies discordantly over Late Cretaceous deposits, which are often included within the Madruga Formation. Clastic material derived from older Cretaceous volcanic-sedimentary rocks are conglomerates that reach from 30 to 40 cm up to 1 to 2 m in diameter. Radiolarian sandstones have been reported (Cushman and Bermúdez, 1948a:68), but the origin of the silica (biogenic or volcanic) is unknown. The age of the unit was originally considered Late Cretaceous by Lewis (1932a:539, 1932b) but later defined as late Paleocene based on several index Foraminifera (Bermúdez, 1950, 1961).

Stratigraphically, the Madruga Formation lies concordant over the Mercedes Formation and discordant over the Peñalver (Late Cretaceous/early Paleocene) and Via Blanca Formations (Late Cretaceous) (Fig. 2). It is covered concordantly by the Capdevilla Formation (early Eocene) and discordantly by the Cojimar Formation (early – middle Miocene) and Nazareno Formation (late middle Eocene – late Eocene). The Cretaceous rocks (K_2_^cp-m^?) can be confused with the overlaying Paleocene layers (Albear et al., 1985; Franco-Álvarez et al., 1992). At the type locality, contacts with other formations are not exposed within the outcrop but are partially exposed at nearby locations. Fossil specimens are sparse and rare throughout the outcrop.

**Figure 2:**
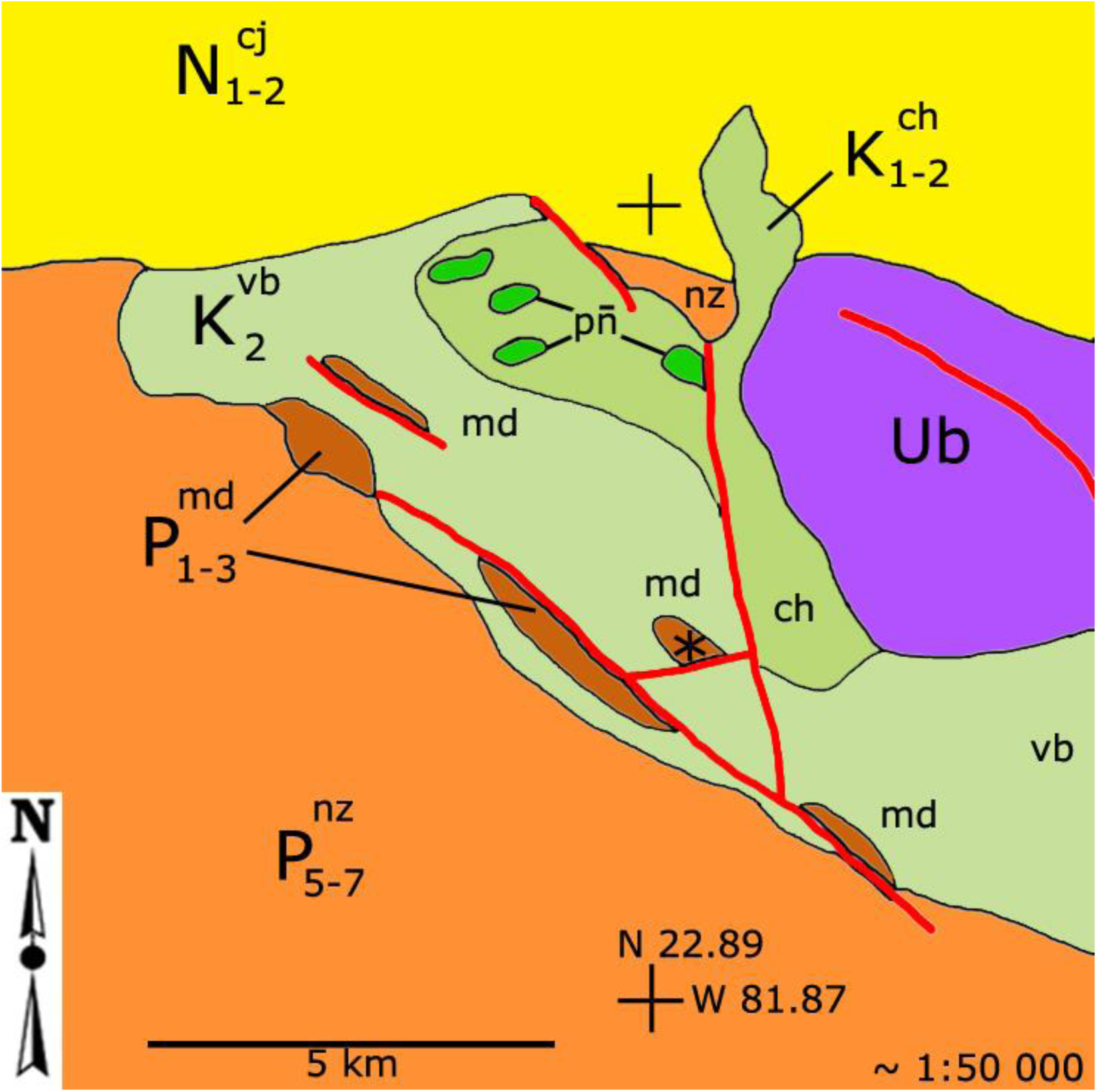
Geological map indicating the outcrops associated with the Madruga Formation at the collection locality discussed (marked with an asterisk *). Madruga Formation (md), Paleocene (P_1-3_), Nazareno Formation (nz) lower Eocene (P_4-5_), Peñalver Formation (pñ) Upper Cretaceous (KTB/K_1-5_), Via Blanca Formation (vb), Upper Cretaceous (Campanian-Maastrichtian, K_5-6_), Cojimar Formation (cjr) lower Miocene (N_1-2_), and Güines Formation (gn) middle-upper Miocene (N_3-4_). UB indicates ultrabasic ophiolite outcrops.

The environment of deposition has been interpreted as a marine synorogenic piggyback basin of medium to deep depth (bathyal), formed under unstable tectonic environment, transportation, and turbidity currents (Bralower and Iturralde-Vinent, 1997; Iturralde-Vinent and MacPhee, 1999; Iturralde-Vinent et al., 2016; Núñez-Cambra and Iturralde-Vinent, 2016).

## RESULTS

### Systematic Paleontology

Class: Bivalvia Linnaeus, 1758

Order: Cardiida Férussac, 1822

Family: Cardiidae Lamarck, 1809

*Cardiinae* Schneider, 2002

Genus: *Schedocardia* Stewart, 1930

*Schedocardia* cf. *hatchetigbeensis* (Aldrich, 1886)

### Specimens

MPAL-1001 (Fig. 3), deposited at the paleontology collection of the Museum of Madruga city. Collected by Yasmani Ceballos Izquierdo on June 2017, at 3-4 m west of the railroad bridge, on the north margin of the roadcut (Figs. 1-2).

**Figure 3:**
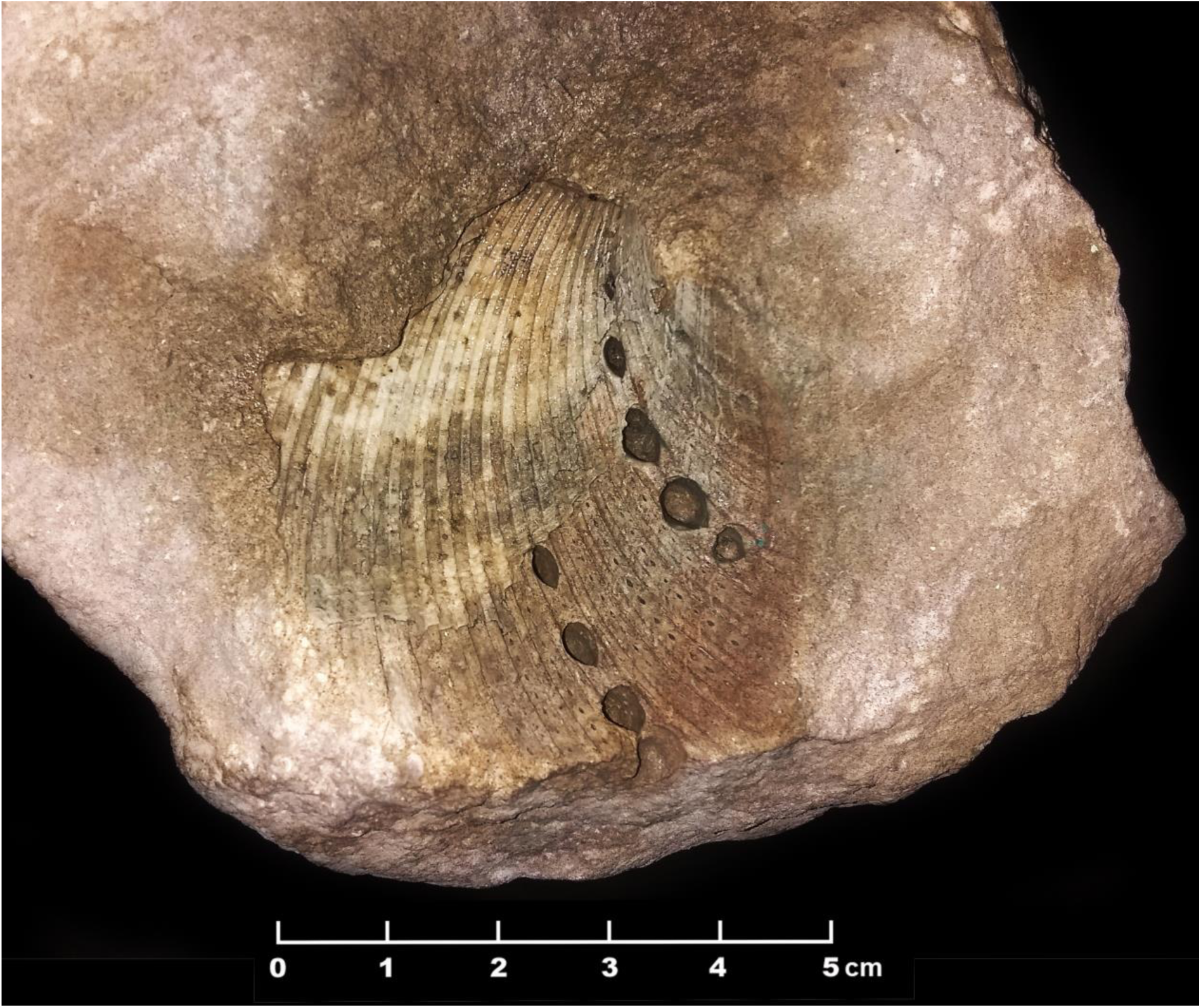
*Schedocardia cf. hatchetigbeensis* (MPAL-1001) right valve, incomplete external mold in the matrix, collected from the Madruga Formation.

### Description

The fossil is an incomplete external mold of the right valve in a calcareous matrix. The external mold is obliquely quadrate, elliptic-ovoid, or schediform in shape (D in Schneider, 2002); radial ribs are narrow and nearly symmetrical throughout. There are small-simple foramina on the interstices between the radial ribs, valve margins crenulated. The interstices between radial ribs are wider than the width of the rib (Fig. 4). External ribs strongly expressed, but not alternating (i.e., wide to narrow) on a central slope. There are >19 ribs present on our specimen, but the exact number is unknown due to incompleteness. The spines or their foramina are more pronounced in the lower valve margins. Only the upper part of the valve external mold is visible. No spines or nobs present. Flanking ribs closely packed together. Cardinals, muscle scars, or hinge not visible (Fig. 4). The umbo is generally incomplete, but likely opisthogyrate and wide. Marked, ovoid-shaped and incised foramina occur almost symmetrically along the curve-space between the valve ribs (Fig. 4). In cross-section, these either extend straight down or curved at the deepest tip. These appear to represent scars of spines that have broken off.

**Figure 4:**
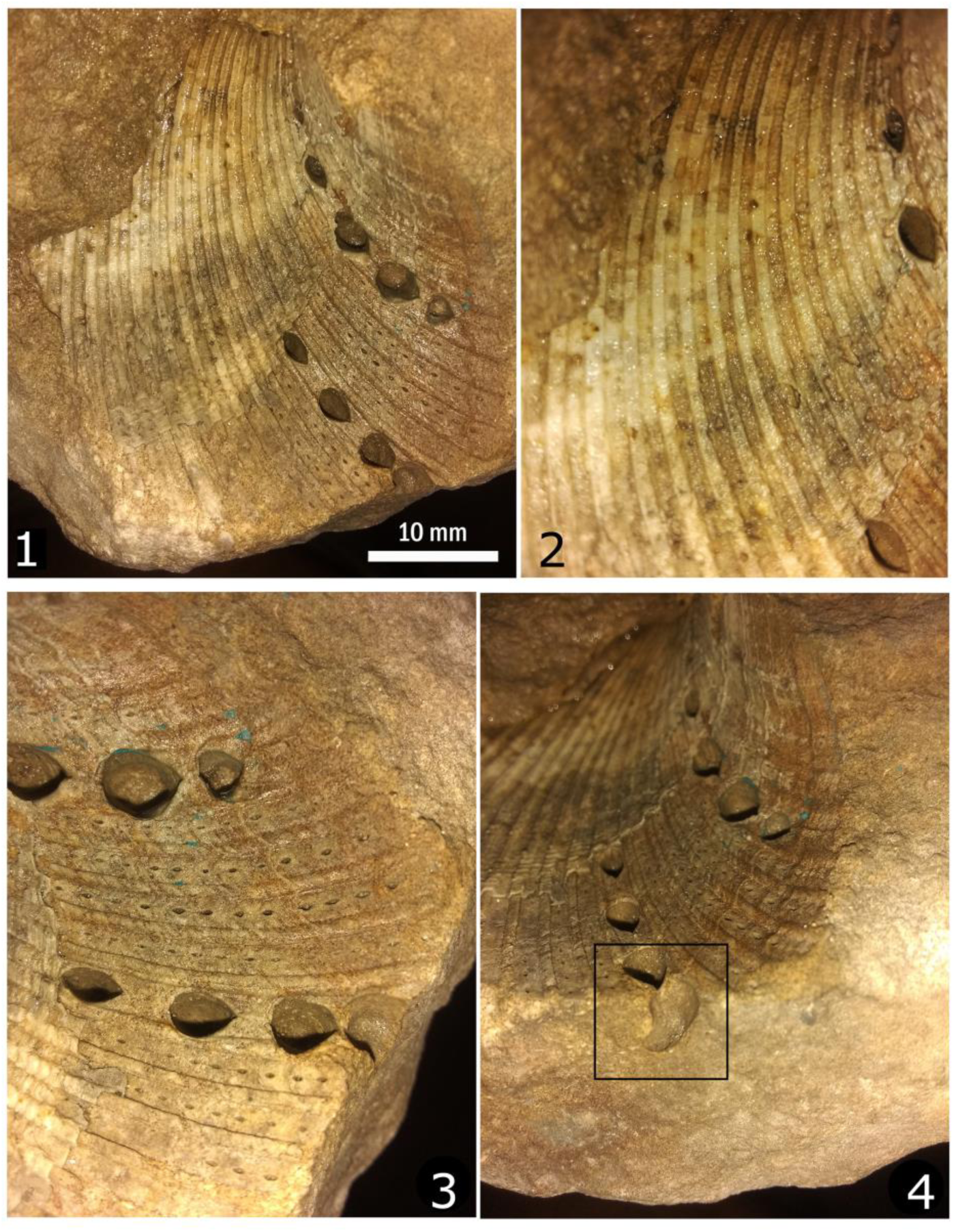
Close-ups of the *Schedocardia* cf. *hatchetigbeensis* (MPAL-1001). Note the ovoid foramen on the valve, and the smaller foramina on the rib tops, suggesting alternating rows of spines.

A smaller cardiid specimen is embedded in the same sample opposite this cast (not illustrated here).

## DISCUSSION

### Identification

The specimen (MPAL-1001) is comparable to *Schedocardia juncea* (Olsson, 1930), and *Schedocardia gatunense* (Dall, 1900), in their elliptic-ovate shape of the valve and higher, less oblique shell anteriorly (Figs. 3-5). However, it differs from these three species in having prominent ribs near the umbo, many more costae, interstices that are flat but wide, not narrow, and ribs that do not bear beads (as in *S. gatunense*). It also differs from *Schedocardia brewerii* Gabb, 1864 in having large spines (Stewart, 1930: 257) more closely packed costae and interstices, and more ovate-elliptic shape of the valve (more elongated in *S. brewerii*, see pl. 3, fig. 11 in Moore, 2002).

**Figure 5:**
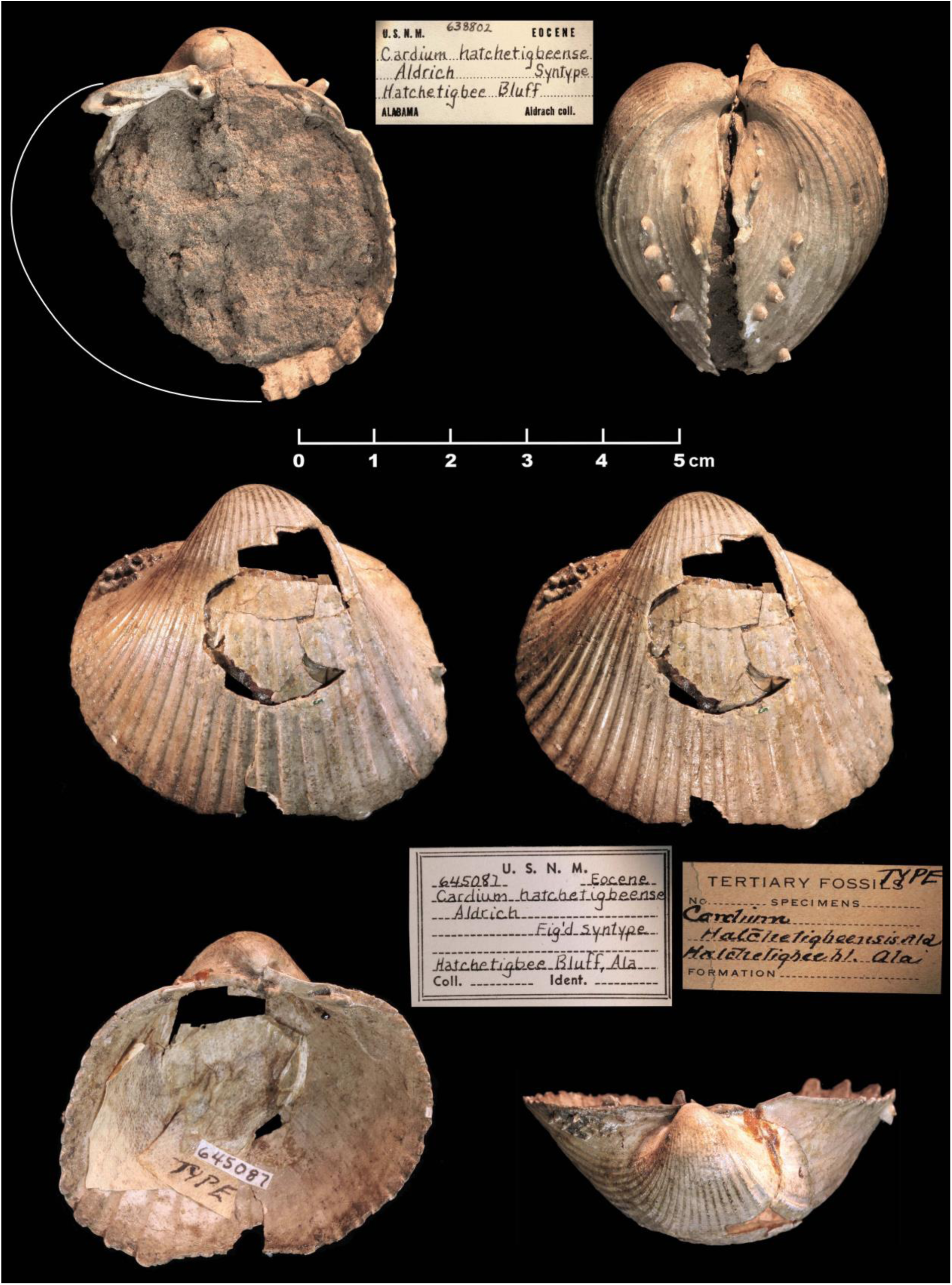
*Schedocardia hatchetigbeensis* syntypes (USNM 638802 =*Cardium hatchetigbeense* and USNM 645087; type =*Cardium hatchetigbeense*) from the Hatchetigbee Formation at Hatchetigbee Bluff, Tombigbee River, Washington County, Alabama. Two specimens in the center row are stereopairs. Photographs used with permission of Jan Johan ter Poorten.

Compared to images of the syntypes of S. *hatchetigbeensis* (Aldrich, 1886) (USNM 638802 and 645087, both from Hatchetigbee Bluff, Alabama, USA) the specimen from Madruga Formation agrees closely with this species in having several radial rows of tiny scars alternating with larger scars, narrower interstices, lacking rib beads, low spines inside these spaces, and ovate shape of the valve. However, the species is not fully identical, and may represent a new form. Due to their overall similarity, we tentatively refer it here to *S.* cf. *hatchetigbeensis* until further specimens are available (Figs. 5-6).

**Figure 6.**
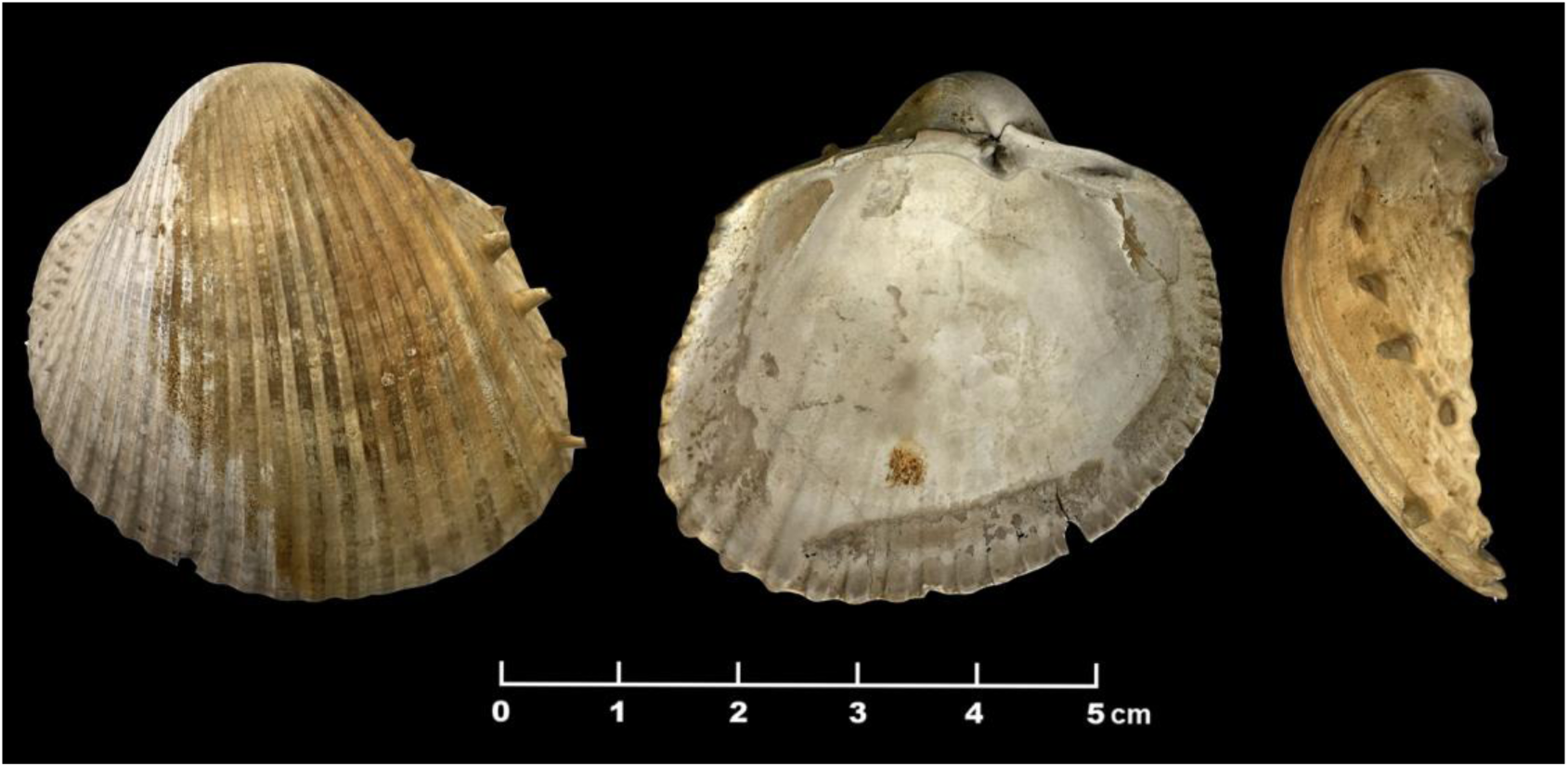
Left valve of *Schedocardia hatchetigbeensis.* Specimen image courtesy of Geological Survey of Alabama. Specimen catalog number GSA-I8608 labelled as *Acanthocardia hatchetigbeensis* from the Hatchetigbee Formation at Hatchetigbee Bluff, Tombigbee River, Washington County, Alabama,

In addition to the *Schedocardia*, another cardiid bivalve, *Acanthocardia* (MPAL-1002), was discovered in the same deposit (Fig. 7). This specimen has an obliquely quadrate shell, with 21 visible ribs, crenulate valve margins, and spinose/nodose ribs. Probable double ribs can be seen on the imbricated valve side. Although the specimen is incomplete, it resembles in overall morphology *Acanthocardia tuberculata* (Linnaeus, 1758) and *Acanthocardia echinata* (Linnaeus, 1758). However, the specimen is too incomplete to assign to species.

**Figure 7:**
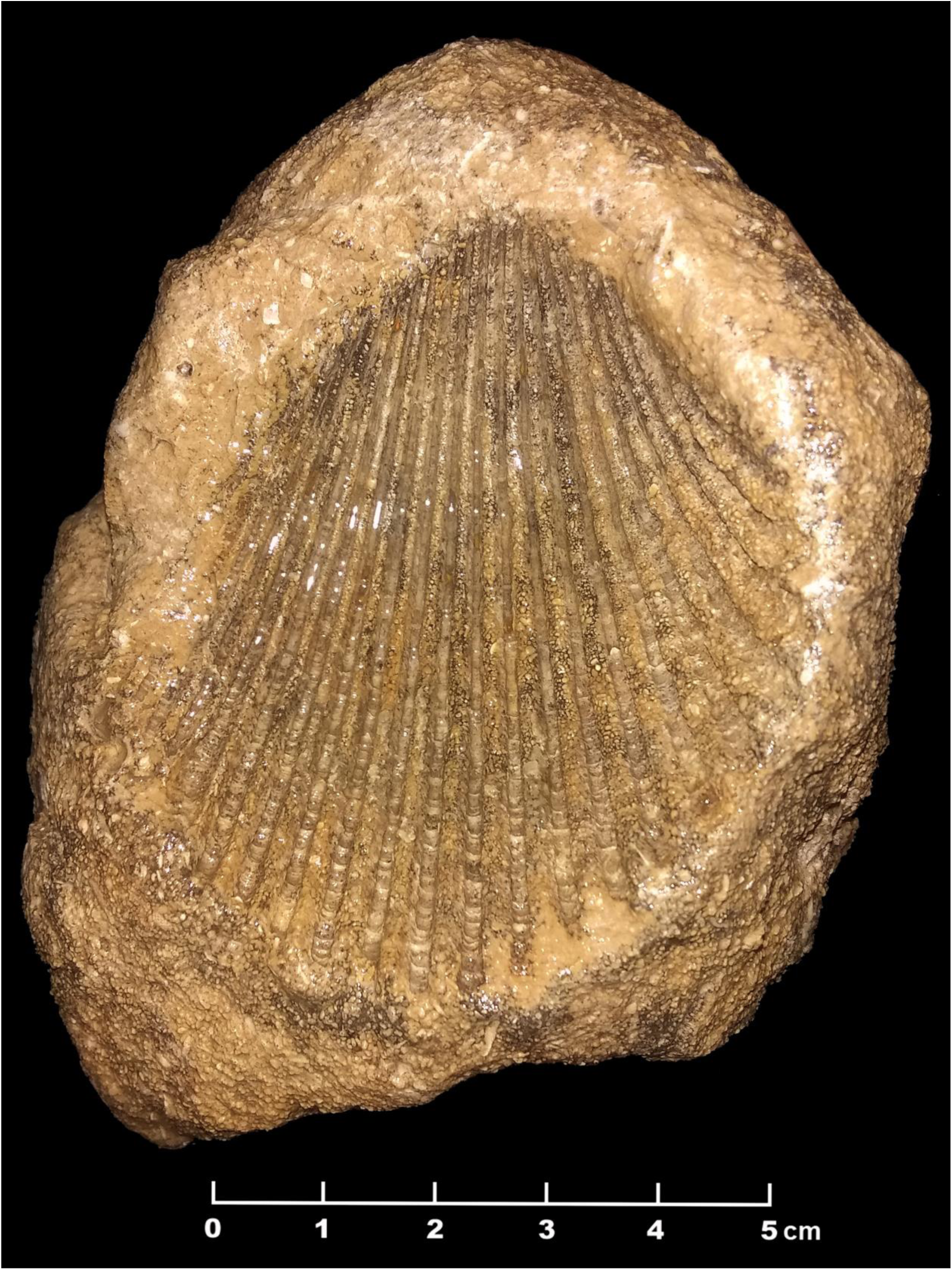
*Acanthocardia* sp. (MPAL-1002) external mold in matrix, collected from the indicated outcrop of the Madruga Formation.

### Chronology

The Madruga Formation has been formerly assigned to the late Paleocene (upper) by the researchers of the Cuban Geological Lexicon and others (Sánchez-Roig, 1949; Sachs, 1957; Bermúdez, 1950, 1961; Furrazola-Bermúdez et al., 1964; Franco-Álvarez et al., 1992) based on the presence of several index taxa, such as the planktic foram *Morozovella velascoensis* (Cushman, 1925), index of PKF zone P5, and *Pseudophragmina* (*=Athecocyclina*) *stephensoni* (Vaughan, 1929), a large benthic foraminifer of carbonate banks (Özcan et al., 2019). More recently, researchers have extended the age of the formation to the early Eocene (Ypresian) due to the presence of the LBF *Euconuloides wellsi* (Cole and Bermúdez, 1944), which is absent in Paleogene assemblages (Blanco-Bustamante et al., 1999).

The genus *Schedocardia* appears in the fossil record in the late Paleocene (Keen, 1980; Schneider, 2002), but did not have its acme until the Eocene. *Schedocardia hatchetigbeensis* has been reported from the early Eocene (Ypresian) of Alabama and Texas, USA (Toulmin, 1977; Garvie, 2013; Sessa et al., 2012). Other *Schedocardia* species, such as *S. juncea* and *S. gatunense* are reported from the late Eocene of Panama, Venezuela, Colombia, and Peru (Woodring, 1982). Therefore, the genus is generally considered to be an Eocene indicator (Woodring, 1982:542). The occurrence of *Schedocardia* in the Madruga Formation seems to support the extension of the Cuban formation at least to the early Eocene. Cushman and Bermúdez (1948a, b) correlated the Madruga Formation with late Paleocene formations from Alabama and Texas, so it is not so surprising that *S.* cf. *hatchetigbeensis* appears here. These North American formations are now considered early Eocene (Toulmin, 1977; Sessa et al., 2012).

### Environment

*Schedocardia* and *Acanthocardia* are considered facultative mobile infaunal (benthic) suspension feeders (Clarkson, 1986). Their occurrence, along with multiple large benthic forams (LBF’s), echinoids, and brachiopods suggests that after the late Paleocene, during the early Eocene, the area of deposition became a shallower marine environment, or were redeposited from shallower environments into deeper by more turbid environments (Blanco-Bustamante et al., 1999). The presence of the conglomerates and polymictic elements may suggest closer proximity to a deltaic environment. Moreover, these suggest that at least some of the fauna reported for the Madruga Formation may have originated in shallower waters of a nearby shelf-forereef environment, transported or reworked by currents or other agents into deeper depositional environments interpreted in the current literature (Albear et al., 1985; Franco-Álvarez et al., 1992; Iturralde-Vinent, 2011a). This interpretation agrees with the overall trend in the orogenic activity of the Cuban terrain during the Paleocene-early/middle Eocene (Bralower and Iturralde-Vinent, 1997; Iturralde-Vinent and MacPhee, 1999; Iturralde-Vinent, 1995, 2011a, b; Iturralde-Vinent et al., 2016).

### Limitations and final considerations

Our observations call attention to several limitations that must be considered for our current interpretation of this outcrop and its fauna. The construction of the highway highly disturbed the area of the outcrop. At the time of the Palmer and Bermúdez visits (the 1930’s-1940’s), those outcrops were well-exposed due to the recent highway construction but are currently severely deteriorated or disturbed by urbanization. Windows into the fossil-bearing strata are rare due to vegetation and soil cover (YCI unp. observations). Former research, however, did not take into consideration the anthropogenic alterations inflicted on the local stratigraphy by the construction of the roads (Figs. 1, 8, and 9). These modifications seem to have severely altered or mixed the external bedding at our collecting locality (Fig. 9), and the horizontality of several beds at several locations. The use of dynamite and heavy machinery likely affected the local stratigraphy and occurrence of its fossil fauna, which up to now has not been considered in any of the published accounts of the outcrop since the 1920’s (Figs. 8 and 9). These human-caused alterations could explain some of the intermixing of the pale Late Cretaceous clastics within some parts of the brown Madruga Formation.

**Figure 8:**
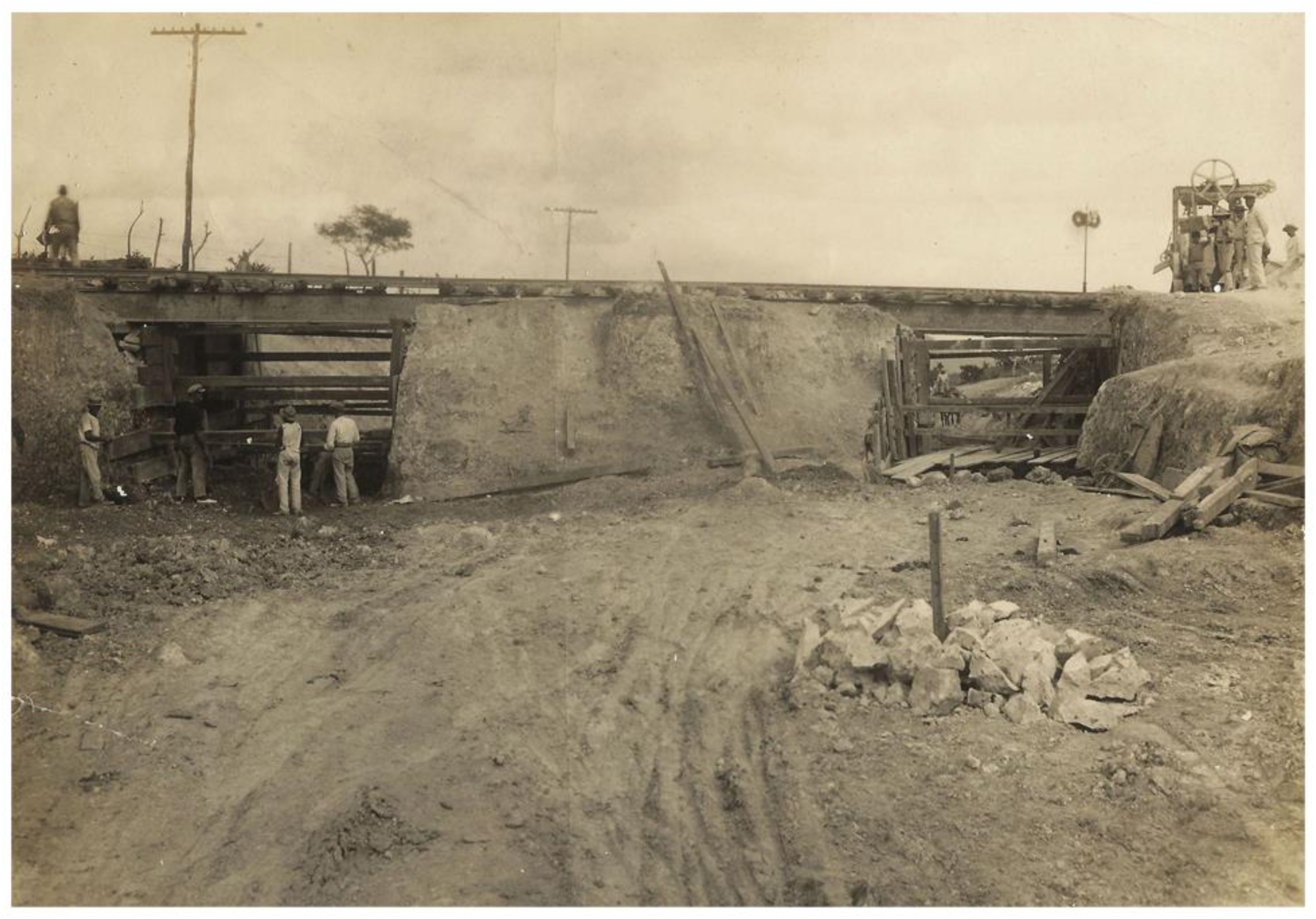
Historical photograph of the road under the railroad bridge during construction in 1928.

**Figure 9:**
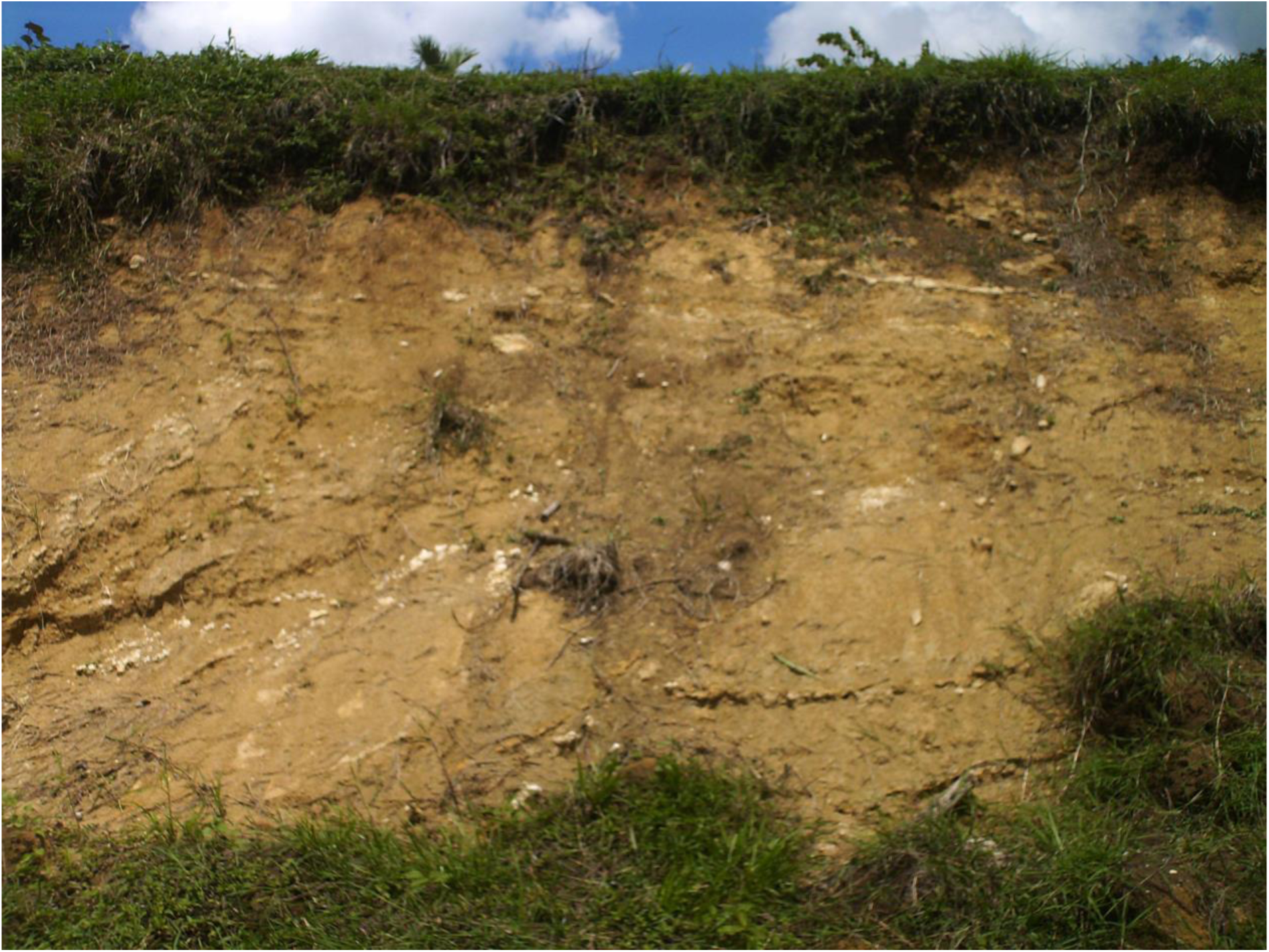
Modern photograph of a section of the Madruga Formation outcrop, at the type locality, in which the exposed stratigraphic column is visible, along with the effect of the use of explosives on the natural stratigraphy. The specimens discussed here were not collected at this precise location.

Furthermore, the historic photographs taken during the construction of the highway and overpass show the presence of a more lithified outcrop-base than the more friable, marly, polymictic sandstones covered by soil and vegetation of today. It is possible that large boulders now present in the outcrop were positioned there then. Even with all the external modifications caused by the building of the central highway on the Madruga Formation type locality, the mollusk specimens here described are referable to that formation. All these factors, and the presence of these hitherto unreported faunal components prompt a reassessment of the whole outcrop that could help clarify the stratigraphy, fauna, and age of the Madruga Formation and its formational history. Much more data is needed to test some of these initial observations and hypotheses. However, we are confident that such an investigation will reveal a far richer Madruga Formation than currently characterized.

## CONCLUSIONS

The Paleocene bivalve fauna of Cuba has been scantly investigated, although outcrops of marine sedimentary rocks of this age are widespread throughout the island. One of the foremost Paleocene Cuban geologic units is the Madruga Formation, which occurs near the town of Madruga, in Mayabeque Province. This formation has produced diverse foraminifera, brachiopods, echinoids, and bivalves. This last group which up to now had not been reported. Thus, by recording the cardiid bivalve genera *Schedocardia* and *Acanthocardia*, we provide a first report for the Cuban fossil record. Historical insight gleaned from photographs taken during the construction of the highway call attention to the high level of disturbance and anthropogenization which the outcrop has been subjected since the late 1920’s and cautions as to the depositional and chronologic interpretation of the samples taken there since. Moreover, our observations could call for a reassessment of the stratotype outcrop of the Madruga Formation.

## Abbreviations

*Institutional* - Specimen repository: Paleontology collection of the Madruga Museum (MPAL), Mayabeque Province, Cuba. LBF-large benthic foraminifera; PKF, planktonic foraminifera; SBF, small benthic Foraminifera. United States National Museum (USNM), Washington, D.C.

## ACKNOWLEDGMENTS

We thank Manuel Iturralde-Vinent and Lazaro Viñola López (Florida Museum of Natural History) for their observations and comments that greatly improved earlier versions of the manuscript. Jan Johan ter Poorten (Naturalis Biodiversity Center) provided further corrections and suggestions, aided in the identification of *Schedocardia*, and kindly allowed us to use his photographs of USNM. Madruga city historian Carlos Miguel Suárez Sardiñas kindly provided important historical insight, historical photographs, and aid during the fieldwork. Sandy Ebersole and Lynn Harrell (both Geological Survey of Alabama) kindly provided images of the *S. hatchetigbeensis* valve in Figure 6.

